# Protecting Allogeneic Pancreatic Islet Grafts by Engineered Veto

**DOI:** 10.1101/2024.01.05.574389

**Authors:** Yan Qi, Xianghua Zhang, Paula Konigsberg, Janae Cull, Patrick McCoy, Coral Cruz, Kevin Dahl, Uwe Staerz

## Abstract

In today’s clinical practice, general immune suppression regimens are used to prevent transplant rejection. Though highly effective, they impair a patient’s protection against infectious challenges. Strategies are being sought that prevent graft rejection without inhibiting beneficial immune functions. One such approach is based on the classical veto effect that employs donor-derived CD8^+^ T cells to inhibit cellular immune responses. Yet, allogeneic grafts may only be partially protected by classical veto as CD8^+^ T cells may fail to remove organ-specific alloreactive T cells. To induce transplant specific immune unresponsiveness, if not tolerance, it may be necessary to endow grafted tissues with veto. Adenoviral vectors were designed that expressed the CD8 α-chain as transgene and thus conferred the immune inhibitory veto function to cells of grafted tissues. In the present model, adenoviral vector vectors protected transduced allogeneic pancreatic islets from rejection in fully immune competent recipients. These studies demonstrated that a tissue-engineering approached could be used to create universally acceptable transplants.

## Introduction

Survival of allogeneic grafts is presently achieved by the administration of cocktails of immune-suppressive drugs. Although systemic treatment regimens efficiently protect transplants from rejection, they impair the patients’ defenses against infectious agents (1, 2, 3). Improved transplantation protocols would induce specific rather than general immune suppression. Classical veto effect has been suggested as one such approach, in which donor-derived CD8^+^ T cells selectively inhibit cellular immune responses to antigens presented on their surface (4, 5). CD8^+^ T cells delete T cells from the peripheral repertoire in an antigen-specific MHC-restricted manner (6, 7). While activated CD8^+^ T cells represent the most potent veto population, other bone marrow-derived (BM) cells exert veto as well (7, 8). Classical veto results from the unidirectional recognition of veto cells by responding T cells (7,9). The immune-suppressive function of the vetoing T cell is independent of its recognition. It is linked to the surface expression of the CD8 α-chain (CD8). Veto function is lost when CD8 is removed from the cell surface, yet reconstituted by the surface expression of CD8 (10,11).

Veto is best explained by a ‘co-triggering’ hypothesis. T cells that recognize a given cell are ‘vetoed,’ and thus commit suicide if they receive concurrent signals through the T cell receptor (TCR) complex and the α3-domain of the major histocompatibility complex (MHC) class I molecules (12, 13). Thus, the exquisite specificity exhibited by veto is based on the engagement of the TCR. It has been argued that the immune inhibitory activity of classical veto cells could be expanded by the release of TGF-β1 (14, 15). However, CD8 α-chain ‘pseudo-variable,’ once anchored on the cell surface through a hybrid antibody (hAb), efficiently triggers veto (16, 17). Different animal models show that infusion of donor-derived classical veto cells prolongs survival of fully allogeneic grafts (18,19,20). Nevertheless, it has been assumed that owing to its unique specificity, veto induced by the injection of classical veto cells such as activated CD8+ T cells cannot provide complete immune unresponsiveness to non-T cell grafts. That is to say, allo-reactive T cells with tissue-specificity for the transplanted tissue may not be inhibited. It may therefore be necessary to transform grafts into ‘veto’-ing tissues.

In initial feasibility studies, we established that the transfer of CD8 with the help of hybrid antibodies (hAbs) imparted veto to cells of diverse phenotypes (16). Hybrid antibodies are cleared from the cell surface quite rapidly. As the immune system takes several days to raise allo-reactive T cell responses in vivo, hAbs were not suitable for organ transplantation. We explored adenoviral gene transfer vectors to efficiently infect both dividing and non-dividing cells of most phenotypes and induce expression of a transgene promptly (reviewed in 21). Since adenoviral vectors rarely integrate into the host cell genome, they mediate transient gene expression. For our application, this feature may be advantageous. In case a CD8-expressing veto vector (VV) were to infect an endogenous antigen presenting cell (APC), resulting aberrant immune suppressions would self-terminate rapidly.

Here, we describe how VVs can be used to transform cells into immune inhibitory veto cells. We show how they imprint highly specific yet stable unresponsiveness to allogeneic pancreatic islets in fully immune-competent animals. Thus, we present a veto-based tissue-engineering strategy that allows the production of universally acceptable transplants.

## Results

### Veto Vector Design

To develop a tissue-engineering strategy to prevent the rejection of grafts, we first constructed a plasmid-based expression vector. This vector carried the mouse CD8 α-chain as a transgene under the control of a CMV immediate early promotor/enhancers (pCMV-mCD8). For organ transplantation studies, adenoviral VVs were used that were replication deficient due to the deletion of the E1 region (ΔE1) (22). A deletion of the E3 region avoided the down-regulation of MHC class I molecules on transduced cells (23), because veto depends on the engagement of MHC class I molecules by the TCR (12). Both the mouse and the human CD8 α-chains were incorporated into adenoviral VVs (mAdCD8 and hAdCD8). Two adenoviral control vectors were produced, in which CD8 was exchanged for a β-galactosidase gene (AdLacZ) or completely deleted (Ad(empty)).

### Ability of VVs to inhibit T cell responses

Initial studies established that CD8 expression transformed cells of different phenotypes into veto cells. C57Bl/6-(H-2b)-derived fibroblasts (MC57) and lymphoma cells (EL4) were transfected with the plasmid-based VV, pCMVmCD8 (MC-CD8 and EL-CD8), and selected for stable surface expression of CD8 (Figure 1a). They were tested for their ability to inhibit the induction of T lymphocytes in vitro. Mixed lymphocyte cultures (MLC), Balb/c anti-C57Bl/6 (H-2d anti-H-2b), were set up in the presence of CD8-expressing fibroblasts or lymphoma cells. Whereas cytotoxic T lymphocytes (CTL) were activated in the presence of non-transfected fibroblasts or lymphoma cells, the addition of CD8-expressing cells drastically reduced their responses (Figure 1b). Having established their immune-suppressive potential in vitro, we probed whether CD8^+^ MC57 could inhibit allogeneic immune response in vivo.

**Figure 1.**
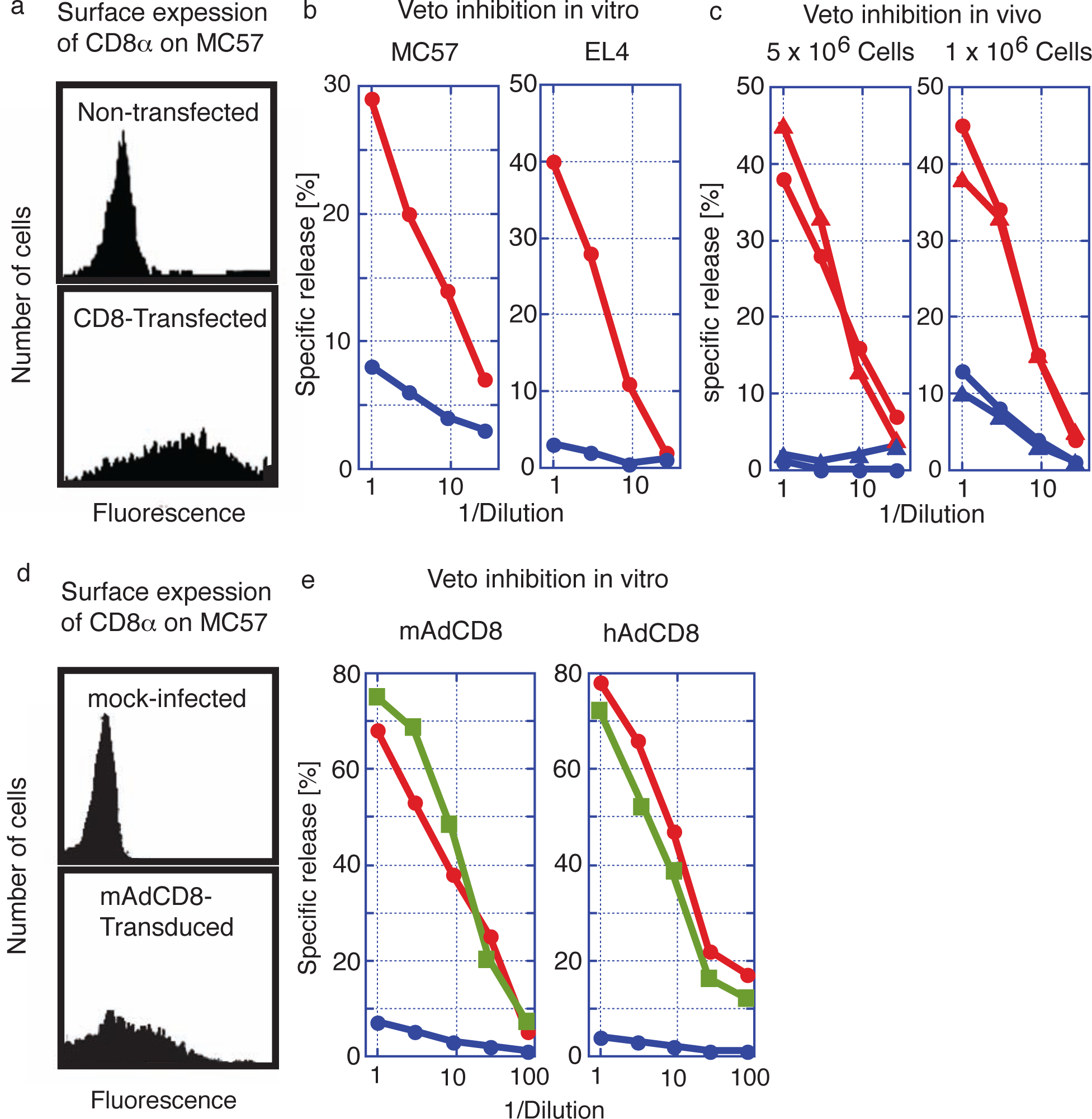

Balb/c (H-2d) mice were injected i.v. with 5 x 10^6^ or 1 x 10^6^ MC-CD8 cells. After three weeks, spleen cells were harvested, challenged in vitro with C57Bl/6 stimulator cells,and tested for the presence of specific CTLs. These studies revealed that the injection of MC57 control cells did not affect the induction of CTLs. In contrast, infusion of MC-CD8 severely limited (1 x 10^6^) or deleted (5 x 10^6^) the activity of H-2b-specific CTLs retrieved from these mice (Figure 1c). These findings proved that cells other than BM-derived cells acquired veto when transfected with a plasmid-based VV.

Plasmids are ill-suited to deliver transgenes to tissues; a more efficient gene transfer vehicle was needed. We therefore evaluated the inhibitory function of the adenoviral VVs, mAdCD8 and hAdCD8, which induced the expression of the CD8 α-chain on approximately 50% of MC57 (Figure 1d). We examined their inhibitory potential in Balb/c anti-C57Bl/6 MLCs, to which control MC57 cells, mAdCD8-, or hAdCD8-transduced counterparts were added. As exemplified in Figure 1e, MLCs supplied with control cells harbored CTL activity similar to those seen in unchallenged cultures. In contrast, the addition of MC57 cells expressing either the mouse or the human CD8 α-chain suppressed the induction of CTLs (Figure 1e). These results firmly demonstrated that adenoviral VVs conferred veto.

### Transduction of pancreatic islets with adenoviral VVs

Having established the efficacy of VVs in transforming defined population of cells, we examined whether veto could be applied to a more complex tissue, such as pancreatic islets. Pancreatic islets were purified from C57Bl/6 mice using conventional enzyme digestion methods (24). To optimize transduction protocols for the Adenoviral VV mAdCD8, both infection time and the multiplicity of infection (MOI) were varied. We observed that an extended (18-hr) virus exposure time pared with a relatively low multiplicity of infection resulted in high transduction efficiencies without affecting islet function (data not shown). Under these conditions, expression of CD8 was primarily seen on surface cells of the pancreatic islets (Figure 2a and b). On average 70% of all pancreatic islets showed evidence of visible CD8 expression (Figure 2c). CD8 could not be detected on non-infected islets, or on islets transduced with the control vector AdLacZ (Figure 2a, b and c).

**Figure 2.**
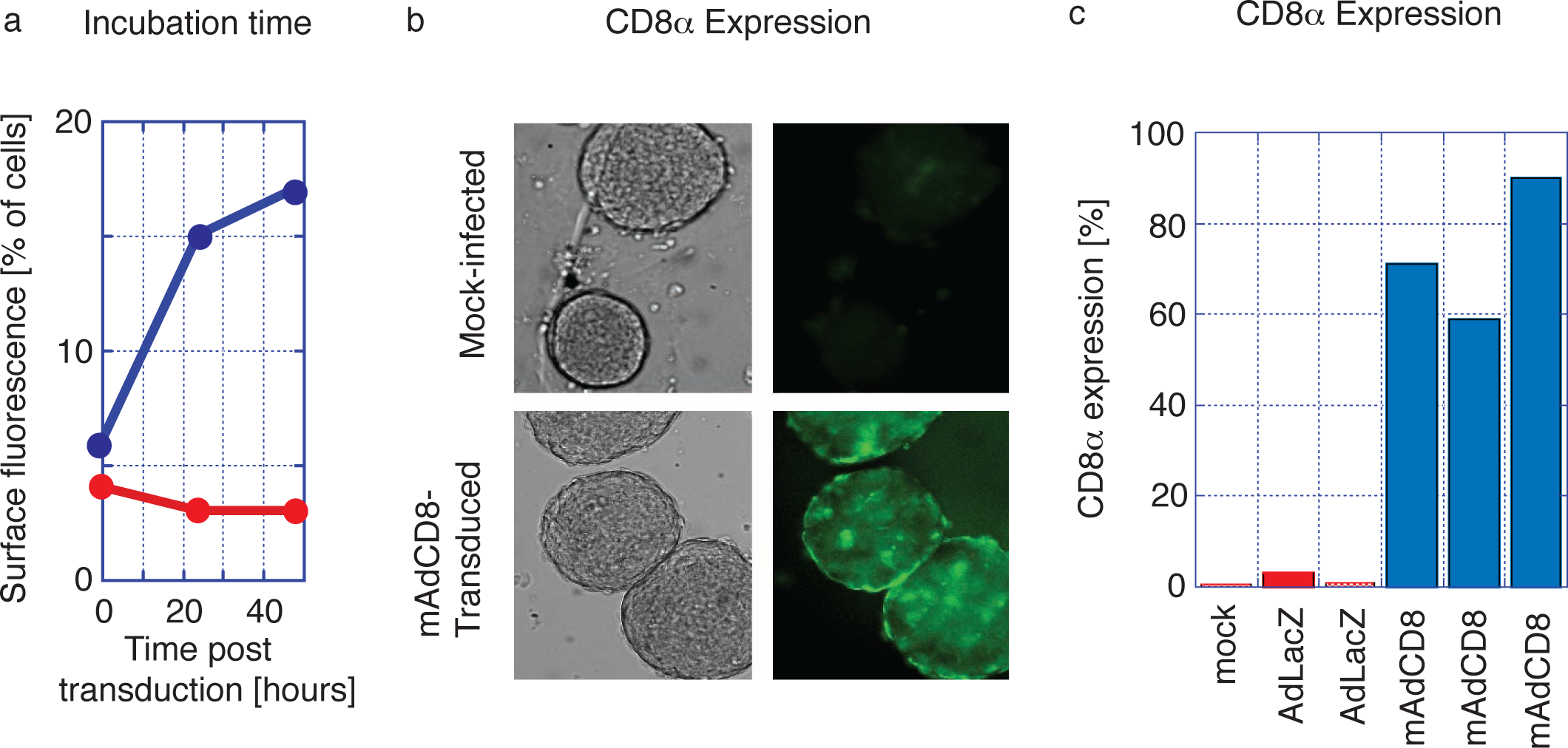

### Protection of VV transduced pancreatic islets from rejection

Having optimized VV transduction protocols, we studied whether VVs would interfere with the rejection of allogeneic pancreatic islets. We chose C57Bl/6 mice as organ donors and Balb/c mice as recipients. These strains are disparate across both MHC class I and MHC class II regions and numerous minor histocompatibility antigens. Therefore, rejection of these allogeneic pancreatic islets should be effective and broad, and as others had shown, mediated by both CD4^+^ and CD8^+^ T lymphocytes (25). Pancreatic islets harvested from C57Bl/6 were either transduced with both mAdCD8 and Ad(empty) or mock-infected, then transplanted under the kidney capsule of Balb/c mice suffering from chemically induced diabetes mellitus. The recipients did not receive any adjuvant immune inhibition and thus remained fully immune competent. In the first set of studies, we increased the number of islets to approximately 800 islet equivalents to compensate for any possible detrimental consequences of the in vitro transduction and to provide more substantial doses of veto cells. As seen in Figure 3a, Balb/c recipients rejected all mock-infected and Ad(empty)-transduced control islets in 23±7 and 28±10 days, respectively. The failing grafts were characterized by cellular infiltration (Figure 3b). In contrast, 83% of mAdCD8-transduced C57Bl/6 islets survived permanently. In these mice the allogeneic pancreatic islets showed no evidence of cellular invasion (as shown in Figure 3b). Immune protection was tested more stringently in another study that used smaller numbers of C57Bl/6 pancreatic islets (450 islet equivalents). As before, mock-infected pancreatic islets were lost swiftly (22±5 days) whereas 91% of mAdCD8-transduced were protected indefinitely from rejection (Figure 3b).

**Figure 3.**
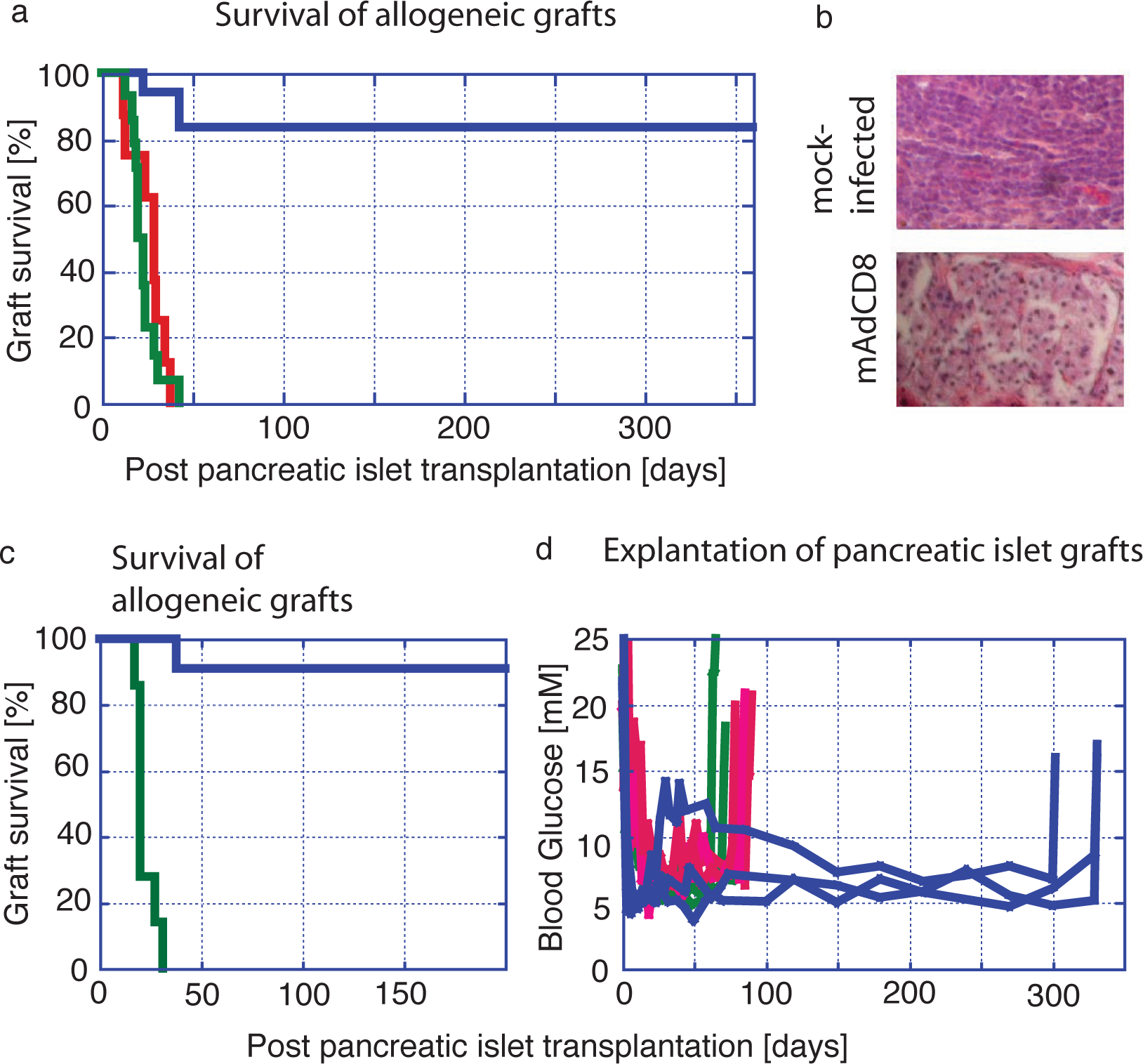

Since control pancreatic islets were destroyed consistently, it was unlikely that a recovery of endogenous insulin production was responsible for the observed normalization of the blood glucose levels. Yet, to formally exclude this possibility, implanted islets were removed by nephrectomy. In mice that had received syngeneic pancreatic islets (C57Bl/6 into C57Bl/6) either VV-transduced or mock-infected, hyperglycemia returned swiftly (Figure 3d). The same held true for Balb/c mice that had carried allogeneic islets under the protection of mAdCD8. Even animals that had maintained allogeneic islets for one year developed diabetes within two days. Therefore, we concluded that pancreatic islet grafts had been solely responsible for the permanent diabetes cures.

### Immune status of transplanted animals

Balb/c mice that had maintained mAdCD8-transduced C57Bl/6 pancreatic islets for at least 6 months were used to investigate how graft acceptance had affected alloreactive responses of both CD8^+^ and CD4^+^ T cells. To evaluate the activity of alloreactive CTLs, spleen cells were challenged in vitro with C57Bl/6 stimulator cells. Balb/c control mice that had not received an islet transplant were tested in parallel. These studies revealed that some of the transplant recipients no longer provided donor-specific CD8^+^ CTLs that could be activated in vitro (Figure 4a). Others retained donor-specific CTLs that nevertheless did not interfere with the presence of the C57Bl/6-derived pancreatic islets.

**Figure 4.**
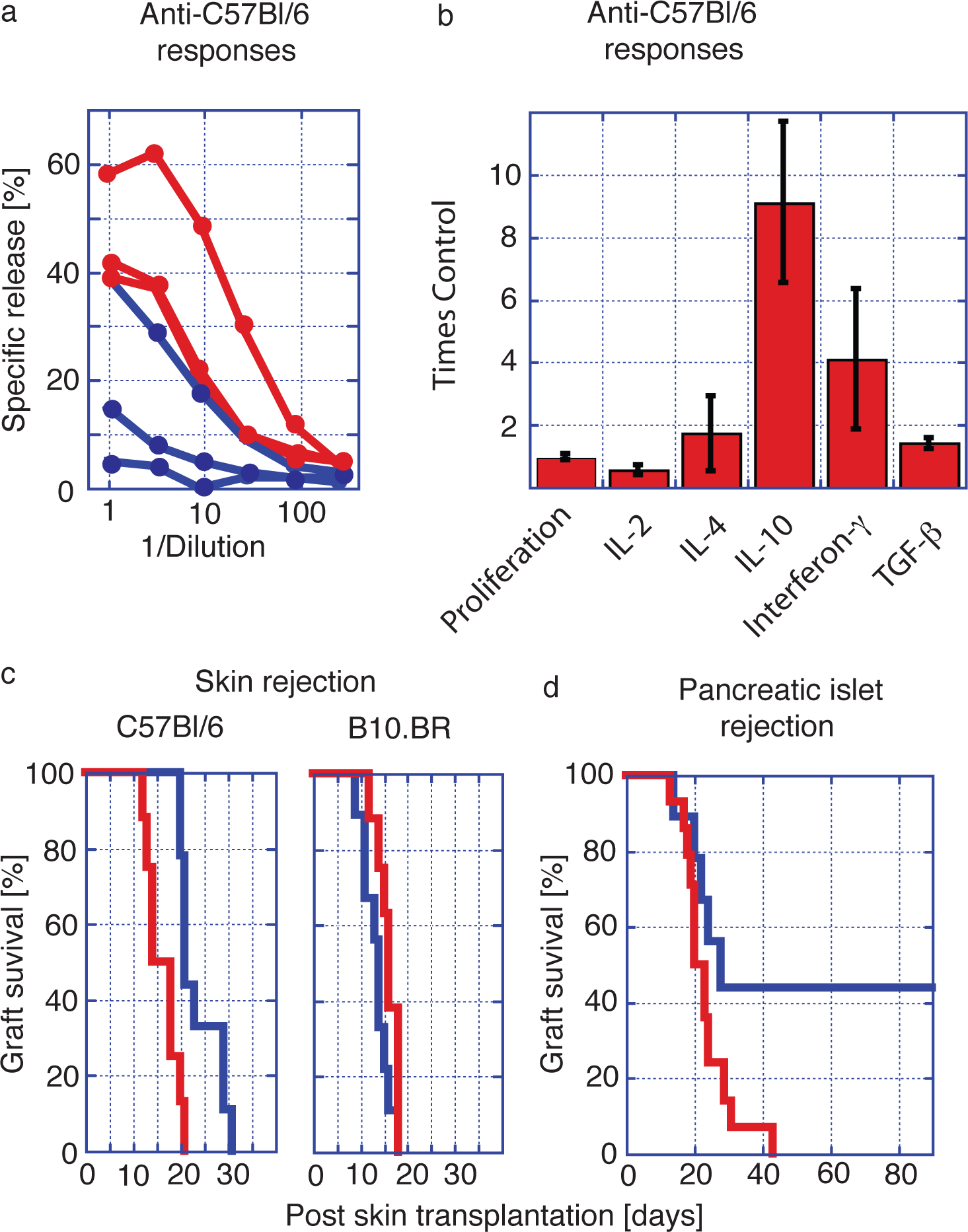

Peripheral CD4^+^ T cells harvested from transplant carriers and control mice were stimulated with Balb/c spleen cells. T cells harvested from control mice and organ carriers proliferated to the same extent. Differences in the release of interleukins (IL) were seen between transplanted and control mice (Figure 4b). A reduction in IL-2 production may indicate a partial inhibition of the activity of alloreactive CD4^+^ T helper-1 cells (26, 27). IL-4 secretion as a marker of T helper-2 cells did not show significant differences. The syntheses of IL-10, interferon-γ (IFN) and TGF-β1 were enhanced in transplanted mice (Figure 1b). This change in the interleukin profile suggested that the presence of the allogeneic graft had expanded the activity of regulatory T cells (28, 29, 30, 31, 32).

We then studied whether the observed immune protection could be broken with a strong immune challenge. Long-term carriers of C57Bl/6 pancreatic islets and Balb/c control mice were transplanted with patches of full-thickness dorsal skins harvested from C57Bl/6 and ‘third-party’ B10.BR donors. We discovered that the rejection of donor-type C57Bl/6 skin was significantly delayed when grafted onto pancreatic islet carriers (Figure 4c). In contrast, B10.BR skins were destroyed swiftly by control mice and longterm carriers of allogeneic islets. This indicated that third-party immune responses had not been affected by the presence of veto-engineered tissue. Thus, VV-transduced grafts had inhibited immune responses in an MHC-haplotype specific manner. Approximately half of all skin-transplanted mice lost their pancreatic islet graft within 30 days (Figure 4d). In these animals allo-reactive T cells had not been completely deleted as they could be activated by a stringent immune challenge. In these mice veto had induced permanent immune unresponsiveness rather than stable tolerance. In a second cohort pancreatic islets remained healthy even though donor-type skin had been rejected (Figure 4d). In these mice, a stable immune tolerance had been imprinted that encompassed broadly reactive and tissue-specific, i.e. islet-specific, alloreactive T cells, and that could not be broken by a most potent challenge afforded by full-thickness skin. Taken together these findings demonstrated that as we had predicted veto-induced immune tolerance was indeed tissue-specific (33, 34, 35).

## Discussion

Even though classical veto had been shown to induce specific immune unresponsiveness, it has not been vigorously pursued for clinical use. Classical vetorequires the enrichment and in the case of CD8+ T cells, the in vitro stimulation of donor derived T cells (36). Infusion of activated donor T cells is not without peril: A graftversus-host disease (GvHD) is set off unless allo-reactive T cells have been removed from the veto cell preparation (37). Although GvHD may be avoided by the use of other BM-derived veto cells, classical veto may nonetheless fail to imprint complete transplantation tolerance. Due to the exquisite specificity of veto, only T cells reactive with antigens presented on veto cells are deleted. Allo-reactive T cells that see grafts in a tissue-specific fashion remain, and may become instrumental in rejecting the transplant (33,34,35). Our studies supported this hypothesis. Even though veto engineered pancreatic islets avoided rejection by the recipient’s immune system, they failed to completely eliminate skin-specific allo-reactive T cells. As a consequence, skin patches harvested from the same donor were not fully protected (Figure 4).

Earlier we had developed hAbs that anchored CD8 α-chain homodimers onto the cell surface (16) and had demonstrated that surface expression of CD8 did transform cells of different phenotypes into specifically immune inhibitory cells (16, 17). These results led us to examine whether complex tissues could be similarly engineered for CD8 mediated veto. Extended CD8 expression was achieved by a switch from the hAb approach to genetic delivery systems. We first used fibroblasts stably transfected with a plasmid-based VV to establish immune inhibitory potential a VV in vitro and in vivo. We switched to a more efficient adenoviral VV for the ultimate transplantation studies. Adenoviral vectors infect most cells, even non-proliferating ones, with high efficiency (21) and thus are well suited to deliver CD8 to a complex tissue. Their transience in expression though detrimental to conventional gene therapy applications, may indeed be beneficial for immune suppression. In the improbable case that host APCs were infected, any aberrant inhibition of the immune system would be self-terminating. In gene therapy, the use of conventional adenoviral gene therapy vectors is also hampered by their strong immunogenicity (38). In our transplantation studies, syngeneic pancreatic islets transduced with adenoviral VV were maintained permanently without any evidence of rejection (Figure 3d). This observation suggested that incorporation of veto into adenoviral vectors lessened their immunogenicity. Indeed, we found in other studies that i.v. or i.m. injection of free adenoviral VVs only raised weak, if any immune responses in mice (data not shown). VV-transduced pancreatic islets successfully evaded immune rejection in fully immune competent allogeneic recipients. These islets were maintained indefinitely in the absence of any adjuvant immune-suppressive therapy.

At the one-year mark, we surveyed grafted islets for the presence of CD8. The results were highly variable, as we found evidence of CD8 expression in only some islets (data not shown). Therefore, the long-term survival of the allogeneic tissue may not depend on the continued presence of CD8. Yet, there seem to be timing and dose effects on the activity of alloreactive T cells. 5 x 10^6^ CD8 expressing fibroblasts functionally deleted CTLs in vivo. 1 x 10^6^ resulted in only a partial inhibition of allo-reactive CTLs. In our pancreatic islet transplantations, the number of CD8^+^ cells transferred to the recipient had to be even lower (Figure 1c). Yet, CD8 expression might have been maintained longer due the slow turnover of pancreatic islet cells.

The small size of the ‘veto-engineered’ tissue might explain the incomplete elimination of allo-reactive T cells seen in some of the animals (Figure 4a, b). Nevertheless, an extended presence of CD8 may have completely suppressed T cell responses in other animals. Our immune response studies suggested that immune regulatory T cells had been induced in mice that maintained pancreatic islets over longer periods of time. In these animals, the permanent protection of veto-engineered pancreatic islets might have been due to the combination of two immune inhibitory effects: the veto deletion mechanism, complemented by the activity of regulatory T cells (30, 32, 39). Even so, specificity of the veto effect was maintained long term.

Our animal studies established the overall feasibility of a CD8-mediated genetic veto engineering strategy. It transformed complex tissues into grafts acceptable to immune competent recipients. Other genetic approaches have been investigated to reduce the immunogenicity of grafts. Since the Fas ligand (FasL) was detected in sites of immune privilege, it had been postulated that it would similarly protect other tissues from immune destruction, as transplants carrying FasL would induce the apoptosis of Fas expressing T cells (40). Transduction studies with the FasL revealed that inflammatory processes were also induced, which resulted in the accelerated rather than delayed destruction of grafted pancreatic islets (41, 42). Additionally, FasL-expressing tissues might inhibit T cells in a non-specific manner, as they may act as a sink for activated T cells, irrespective of their specificity. In another approach, tissues were engineered to secrete immune inhibitory compounds. In the case of CTLAIg, when given systemically or released locally by transduced tissues, it was effective within an immune-suppressive combination therapy (43, 44). TGF-β1 was also successfully used, transforming dendritic cells into highly immune suppressive cells (45). However, a release of TGF-β1 to modify the immunogenicity of other organs would be problematic. For instance, TGF-β1 induces a fibrosis of the liver (46).

In contrast to a release of soluble factors, veto is intrinsically specific, since it relies on the engagement of TCR on engineered tissues. T cells that do not react with the engineered graft will not be affected. We expect that that unique activity of veto can be harnessed for transplantations of many tissues and organs. In preliminary experiments, we have successfully transplanted liver and heart tissues under veto protection (data not shown). We will undertake large animal trials to investigate how adenoviral VV can be optimized for clinical use, by itself or complemented with other immune-suppressive therapies. In addition, we will engineer the VVs as fully deleted adenoviral vectors so avoid the expression of adenoviral genes in the transduced tissues.

## Materials and Methods

### Animals

Male and female mice (C57Bl/6, Balb/c and B10.BR) 8 to 24 weeks of age were purchased from Charles River Laboratory (Boston, MA).

### Expression vectors

pCMV-mCD8 carries the mouse CD8 α-chain driven by the CMV immediate early promotor/enhancer (Invitrogen). The Adenoviral expression vectors, mAdCD8 and hAdCD8, were of human Adenovirus Type 5 and deleted for the E1 and E3 regions (Qbiogen). Expression of the mouse and human CD8 α-chain transgenes was controlled by a CMV immediate early promotor/enhancer.

### Mixed lymphocyte cultures

Spleen and lymph node cells were harvested, and single cell suspensions were prepared. Erythrocytes were lysed with Tris-buffered ammonium chloride (Sigma-Aldrich) and resuspended in tissue culture medium. For the induction of CTLs, unfractionated responder spleen cells (4 x 10^6^ cells/well) were stimulated with irradiated (3,000 rad) spleen cells (2 x 10^6^ cells/well) in flat-bottomed 24-well plates (Becton-Dickinson). The cultures were incubated for four days at 37°C and 7% CO_2_ in a tissue culture incubator (Forma Scientific). Blasting T cells were counted and tested for their ability to specific target cells. CTLs were incubated with 1 x 10^4^ [^51^Cr]-NaCr_2_O_4_-labeled EL4 (C57Bl/6-derived lymphoma) target cells for four hours in round-bottomed microtiter plates (Becton-Dickinson, total volume 0.2 ml tissue culture medium). Varied effector cell numbers were added as indicated in the text. The amount of ^51^Cr released into the supernatant was measured on a Gamma 4000 (Beckman-Coulter). Inhibitor cell populations were added to some cultures at stimulator-to-inhibitor ratios of 3-to-1. They consisted of murine fibroblasts (MC57T) or lymphoma cells (EL4) (both C57Bl/6-derived), mock-infected, stably transfected with pCMV-mCD8, or transduced with mAdCD8 or hAdCD8. Wells containing target cells, but no effector cells were used to determine nonspecific release, and wells containing target cells in the presence of 1% Triton X-100 (Sigma-Aldrich) were used to determine total release. Percent specific release was calculated as: [(cpm released in the presence of effector cells) - (cpm of nonspecific release)] / [(cpm of total release) - (cpm of nonspecific release)] x 100. For measuring responses of CD4^+^ T cells, T cells were enriched on Nylon wool columns. Thereafter, CD8^+^ T cells were removed by chromatography through anti-CD8 affinity columns (Mitenyi). More than 90% of the remaining cells represented CD4^+^ T cells (data not shown). CD4^+^ responder T cells (5 x 10^5^ cells/well) were cultured in triplicate with irradiated (3,000 rad) spleen cells (5 x 10^5^ cells/well) in flat-bottomed 96-well plates (Becton-Dickinson, total volume 0.2 ml of tissue culture medium). The cultures were incubated for four days at 37°C and 7% CO_2_. Supernatant from each culture was taken to determine interleukin levels. Blast cells were counted and then stained for the presence of CD4^+^ and CD8^+^ T cells. Iscove’s modified Dulbecco’s medium (IMDM, Sigma-Aldrich) was used as tissue culture medium, supplemented with Hepes, glutamine, hydroxypyruvate, 2-mercaptoethanol, nonessential amino acids, penicillin, gentamycin (Sigma) and 10% fetal bovine serum (Intergen).

### Flow cytometry

Cells suspended in staining buffer (PBS / 5% fetal bovine serum) were incubated with both phycoerythin-coupled anti-CD4 and fluorescein-isothiocyanate-coupled CD8 (Becton-Dickinson) in 96-well round-bottomed plates (Becton-Dickinson). The unbound antibodies were washed away and the extent of antibodies binding to the cell surface was analyzed on a fluorescence-activated cell sorter (Coulter Epics).

### Cell infusion and tissue transplantation

Balb/c mice were injected i.v. with 1 x 106 or 5 x 106 cells stably transfected with pCMVCD8. Pancreatic islets were harvested from C57Bl/6 and Balb/c mice and transplanted into Balb/c mice, which had developed stable diabetes mellitus induced by the i.v. injection of streptozotocin (Sigma-Aldrich). The preparation of pancreatic islets has been detailed by others (24). After donor mice were euthanized, the peritoneal cavity was opened, and the pancreatic duct identified and cannulated. Digestion was initiated by an infusion of 4 ml of collagenase type 5, dissolved in Hank’s balanced salt solution (Sigma-Aldrich). Digestion continued after the pancreas was placed into a 50 ml tube (Becton-Dickinson) containing collagenase solution (37°C). With vigorous shaking, the tissue digestion continued for about 20 min, at which point the material was strained to remove large tissue fragments. The released pancreatic islets were spun down, re-suspended and purified utilizing a Histopaque (Sigma-Aldrich) gradient. Researcher-determined islet equivalents were subsequently hand-selected under a surgical microscope (Stereomaster, Fisher).

The purified pancreatic islets were incubated in RPMI 1640 / BSA (Sigma-Aldrich in a 15 ml tube (Becton-Dickinson, 500 islets per tube) for 18 hours at 37°C and 7% CO^2^. Some pancreatic islets were transduced with mAdCD8 or Ad(empty). The vectors were added at an approximate multiplicity of infection (MOI) of 3-to-1 (plaque forming units to pancreatic islet surface cells). Upon completion of infection, free virus was washed away, and the pancreatic islets were re-incubated for 24 hours.

Pancreatic islets were stacked by centrifugation into teflon tubing. They were placed into small pouches that had been created under the capsule of the left kidney of recipient mice. C57Bl/6 pancreatic islets (800 islet equivalents) untreated (14 animals), transduced with Ad(empty) (8 animals) or with mAdCD8 (18 animals), were transplanted into Balb/c recipients (Figure 3a). Of the latter group, 9 animals were observed for about 180 days and 9 animals for about 1 year.

In another trial, 450 C57Bl/6 pancreatic islet equivalents untreated (7 animals) or transduced with mAdCD8 (11 animals) were transplanted Balb/c mice (Figure 3c). These animals were studied for 200 days. In some mice, transplanted pancreatic islet implants were removed by nephrectomy of the left kidney (Figure 3d). In all grafted mice, blood glucose levels were determined twice weekly as a measure of pancreatic islet activity by evaluating tail vein blood using a glucometer (TheraSense).

Full-thickness dorsal skin was harvested from C57Bl/6 and B10.BR mice and all fatty tissue was carefully removed. Skin segments (approximately 1 cm x 1 cm in size) had already been removed from the dorsal skin of recipient mice. They were replaced by donor skin of the same size. Nine Balb/c mice that had carried mAdCD8-transduced C57Bl/6 for at least 6 months were transplanted, as well as 9 untreated Balb/c control animals (Figure 4). Viability of the skin transplant was determined visually.

### Histology

Kidneys harvested from pancreatic islet transplanted mice were fixed with 10% neutralized formalin (Sigma-Aldrich), paraffin embedded, sectioned (Vibratom, TechnicalProducts International), and stained with H&E (Sigma-Aldrich).

### Interleukin assays

Supernatants harvested from mixed lymphocyte cultures established from three mice of each group were tested using commercial assay kits (e.Bioscience, R&D Systems, Bender MD Systems) for the presence of interferon-γ, IL-2, IL-4, and IL-10. The amount of cytokines released was calculated with the help of a standard curve of known lymphokine concentrations.

## Notes

### Competing Interest Statement

The authors worked for Greffex, Inc., a biotechnology company, located in Aurora, Colorado, USA. Besides their salary, they were compensated with grants of shares and/or stock options.

